# A versatile microfluidics platform for enhanced multi-target super-resolution microscopy

**DOI:** 10.1101/2025.10.07.680900

**Authors:** Samrat Basak, Kim Chi-Vu, Nikolaos Mougios, Nazar Oleksiievets, Yoav Pollack, Sören Brandenburg, Felipe Opazo, Stephan E. Lehnart, Jörg Enderlein, Roman Tsukanov

## Abstract

DNA-based Point Accumulation for Imaging in Nanoscale Topography (DNA-PAINT) is a powerful variant of single-molecule localization microscopy (SMLM) that overcomes the limitations of photobleaching, offers flexible fluorophore selection, and enables fine control of imaging parameters through tunable on- and off-binding kinetics. Its most distinctive feature is the capacity for multiplexing, which is achieved through a process known as Exchange-PAINT. This technique involves assigning orthogonal DNA strands to different targets within a sample and then sequentially adding and removing complementary imager strands that are specific to only one target at a time. However, manual Exchange-PAINT workflows are often inefficient, prone to drift and variability, and lack reproducibility.

Here, we introduce a custom compressed-air-driven microfluidics system specifically designed for multiplexed SMLM. Featuring a stackable and modular design that is, in principle, not limited by the number of channels, the system ensures robust, reproducible, and material-efficient buffer exchange with minimal dead volume. It operates in both manual and automated modes and can be readily adapted to a wide range of commercial and custom microscopes, including wide-field, confocal, STED, and MINFLUX platforms.

We demonstrate robust 5-plex Exchange-PAINT imaging in cancerous U2OS cells and, importantly, we establish multiplexed nanoscale imaging in fragile primary cardiomyocytes. These applications highlight the unique power of our platform to extend super-resolution multiplexing into physiologically relevant systems, thereby opening new avenues for biomedical research.

## Introduction

Over the last decade, multiplexed super-resolution microscopy (SRM) has undergone tremendous development, providing unprecedented insights into the nanoscale architecture of biological systems. By enabling the simultaneous or sequential visualization of many molecular species, multiplexed SRM facilitates studies of protein colocalization, organelle organization, and molecular interactions in complex biological environments. Achieving nanometer-scale localization precision, however, requires not only photophysical optimization but also precise environmental control within the experimental chamber.

DNA-PAINT (Point Accumulation for Imaging in Nanoscale Topography), introduced by Jungmann and colleagues in 2010^1^, has emerged as one of the most powerful single-molecule localization microscopy (SMLM) methods^2^. Based on transient binding of short fluorescently labeled “imager strands” to complementary “docking strands” conjugated to target molecules, DNA-PAINT achieves virtually unlimited imaging time and high labeling density, free from photobleaching limitations^3,4^. Its most striking feature is the capacity for multiplexing: by assigning unique DNA sequences to different targets, sequential imaging can be achieved via Exchange-PAINT^5^, while parallel multiplexing is enabled by spectral separation^6,7^ or fluorescence-lifetime DNA-PAINT (FL-PAINT)^8^.

Recent methodological advances highlight both the power and challenges of highly multiplexed imaging. The introduction of secondary labels-based methods, SUM-PAINT^9^ and FLASH-PAINT^10^ significantly enhanced the performances (in terms of versatility and imaging speed) of multiplexed DNA-PAINT imaging, enabled neuronal imaging with up to 30 targets^9^. Another versatile multiplexing approach, based on erasable labels, called Nanoplex, pushed multiplexed STED microscopy to 21-color imaging^11^. Most recently, resolution enhancement by sequential imaging (RESI) has set a new benchmark for molecular-resolution mapping of complex protein assemblies^12^. However, all these approaches rely on repeated manual buffer exchanges, which are tedious, time-consuming, and susceptible to variability and sample drift. Microfluidics can directly address these challenges, making Exchange-PAINT-based methods, and related approaches faster, more reproducible, and reagent-efficient, thereby unlocking their full potential for structural biology investigations.

By allowing precise and programmable fluid handling at the microliter scale, microfluidic systems provide precise control over the sample environment, minimize mechanical drifts in optical systems, and enable reproducible solution exchange across many cycles. The pioneering work of Quake and colleagues on pneumatic valve control^13^ laid the foundation for modern microfluidic devices^14,15^, while the seminal contribution of Nir and co-workers demonstrated the integration of microfluidics into DNA nanotechnology^16^, paving the way for automated workflows in DNA-based artificial motors^17^. Microfluidics has since become indispensable across biology, chemistry, and medicine^18^, but its adoption in super-resolution microscopy remains limited^19^. Existing syringe-based microfluidics platforms^20^ are efficient but constrained by bulky hardware and finite channel capacity, posing a limitation for highly multiplexed workflows requiring multiple sequential buffer exchanges. Additional interesting innovation included integration of pipetting robot for the full automation of SRM imaging workflow^21^.

Here, we introduce a custom compressed-air-driven microfluidics platform specifically designed for multiplexed SMLM imaging and compatible with a wide range of SRM modalities. Our system features a modular, stackable design that is in principle not limited by the number of fluidic channels, thereby overcoming scalability issues of syringe-based approaches. It combines rapid, precise, and material-efficient buffer exchange with minimal dead volume, operating in both manual and fully automated modes. Importantly, the system can be seamlessly integrated with custom-built and commercial wide-field, confocal, STED, and even MINFLUX microscopes.

We validate the performance of our system in demanding biological applications, demonstrating five-target Exchange-PAINT imaging in U2OS cancer cells as well as, for the first time, in isolated ventricular cardiomyocytes. The latter showcases the feasibility of multiplexed nanoscale imaging in fragile and complex primary cells, opening new opportunities for advanced biomedical research. Together, these capabilities position our microfluidics platform as a robust and broadly enabling technology for high-throughput, reproducible, and scalable multiplexed super-resolution imaging.

## Results

### Design and operation of the microfluidics system

To overcome the limitations of manual Exchange-PAINT imaging workflow, we developed a compressed-air-driven microfluidics platform tailored for multiplexed SMLM. The system features a modular, stackable design that can, in principle, accommodate an unlimited number of inlet channels. Each module contains individually addressable valves actuated by externally applied air pressure, eliminating the need for bulky syringe pumps or complex actuators. This compact design reduces the system footprint and allows straightforward scaling of channel capacity according to experimental needs.

In operation, compressed air pressurizes reagent tubes, driving solutions into the experimental chamber at controlled flow rates. Only low pressures (<1 bar) are required, making the system compatible with both laboratory air lines and portable compressors. Air is directed through electronically controlled solenoid valves that select the specific channel for solution delivery. Small-volume tubes (2 mL) were typically used for imager strands, while larger tubes (15 mL) supplied wash buffers. To minimize dead volume and prevent gravity-driven backflow, tube holders were mounted close to the sample and aligned at chamber height, see Figure 1A. Solutions were delivered via biocompatible tubing and removed either manually or, for sensitive applications, using a peristaltic pump for gentle and consistent washing, as shown in Figure 1B.

**Figure 1.**
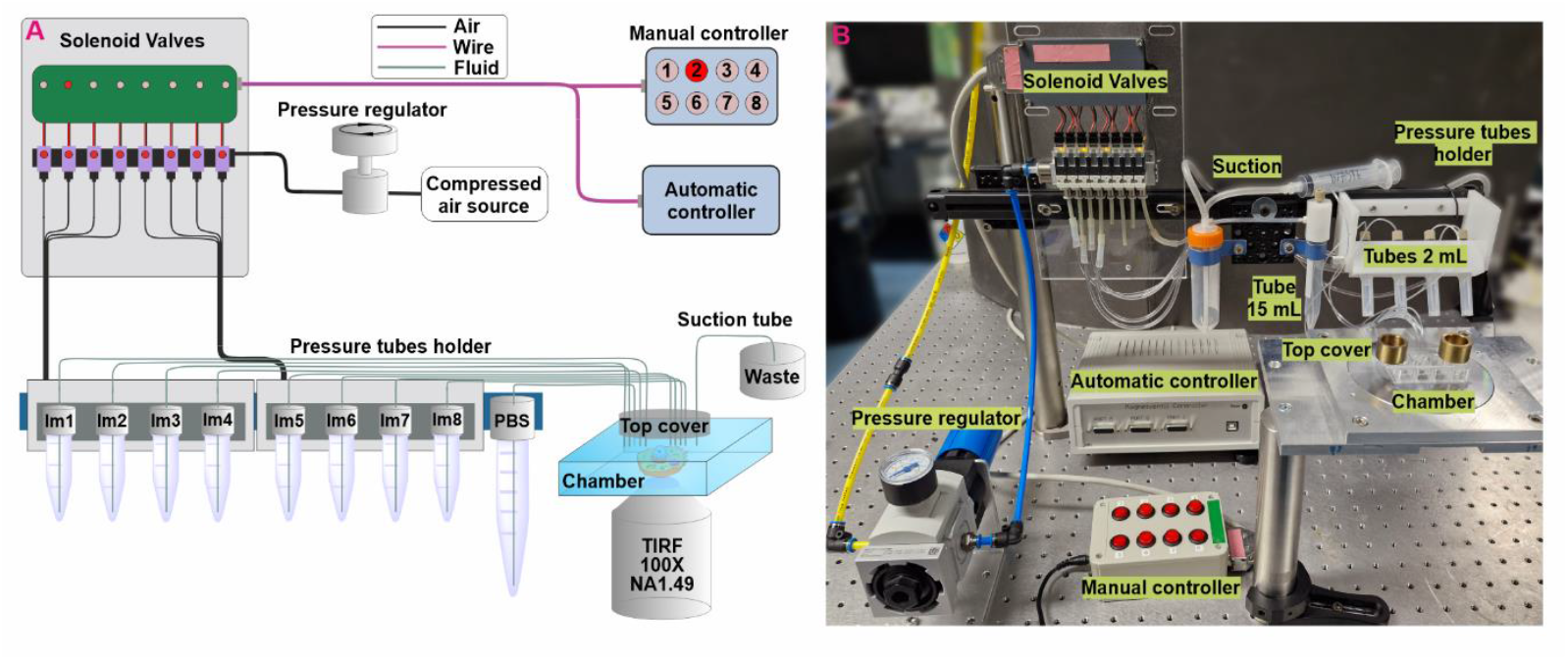
Microfluidics system for multiplexed SMLM imaging. (A) Schematic of the microfluidics platform showing the main components: manual or automated controller, solenoid valve manifold for distribution of air pressure, pressurized reagent tubes, sample chamber with tubing connections, and solution removal syringe. (B) Picture of the assembled microfluidics setup. The modular and stackable architecture enables flexible scaling of channel number, while gentle and reproducible fluid handling ensures compatibility with multiplexed super-resolution imaging workflows.

The system achieves fast and reproducible buffer exchange with minimal reagent consumption. For a commonly used Petri dish, complete exchange was achieved within 30 s, using as little as 1 mL per cycle. The response time between valve actuation and solution arrival at the sample was consistently below 1 s, and carry-over between solutions was negligible, as confirmed by DNA-PAINT experiment.

Control of the system was implemented via either a simple manual switchbox or an automated controller, with support for up to 48 channels, as detailed in Supplementary. Importantly, the open design allows straightforward adaptation to other programming environments such as Python or MATLAB, facilitating integration into custom or evolving laboratory infrastructures. Automated workflows allowed precise programming of sequence, timing, and duration of each solution exchange, ensuring reproducibility and minimizing user-dependent variability.

Importantly, the fluidics system operates independently of the optical path and is broadly compatible with both commercial and custom-built microscopes. We validated its integration with wide-field and confocal SMLM setups, and its design makes it readily deployable across diverse imaging platforms without requiring modifications to the microscope body or detection scheme.

Together, these design features establish a versatile, scalable, and user-friendly microfluidics solution that overcomes the inherent limitations of manual workflows and enables automated, high-precision, and reproducible multiplexed SRM workflows.

### Five-target multiplexed imaging of the cytoskeleton and focal adhesions in U2OS cells

To benchmark the performance of our microfluidics platform, we conducted five-target Exchange-PAINT imaging in U2OS cancer cells. For this test, we selected a panel of proteins at the cytoskeleton–focal adhesion interface, including microtubules, vimentin, and actin as representative cytoskeleton elements, along with the focal adhesion proteins zyxin and paxillin, as shown in the schematics of Figure S1 in the SI. This selection provided a rigorous benchmark, as it required resolving both extended cytoskeletal filaments and nanoscale adhesion scaffolds within the same cell. We used a flexible antibody–nanobody labeling strategy that enabled efficient multiplexing with negligible crosstalk and high labeling efficiency across all channels^22^.

High-quality super-resolution images were obtained for each target (Figure 2, Supplementary Figures S4 and S5), with an average localization precision better than 11 nm (Table S1 in the SI). The structures of microtubules and vimentin filaments were clearly resolved, while focal adhesion proteins zyxin and paxillin displayed distinct nanoscale patterns consistent with their known subcellular distributions^23^. Cross-correlation analysis using Pearson’s coefficient further confirmed these relationships, showing a strong overlap between cytoskeletal structures (e.g., actin with microtubules and vimentin). As expected from their spatial organization, a reduced correlation was observed between the cytoskeleton and focal adhesion proteins.

**Figure 2.**
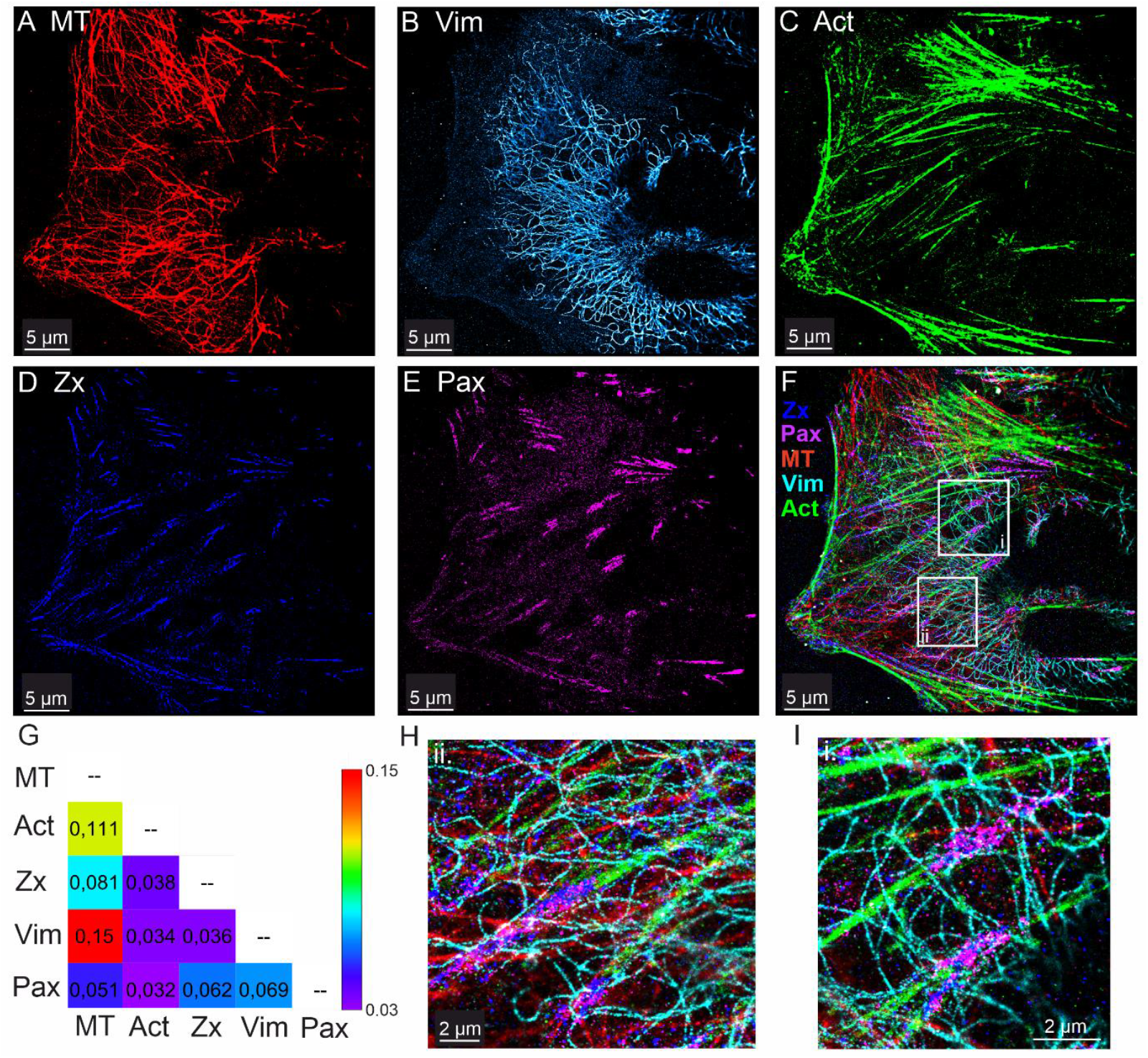
Microfluidics-enhanced Exchange-PAINT 5-target imaging of U2OS cell. Sequential imaging of cytoskeletal and focal adhesion targets: (A) microtubules (MT), (B) vimentin (Vim), (C) actin (Act), (D) zyxin (Zx), (E) paxillin (Pax), and (F) composite overlay of all channels. Scale bar 5 µm. (G) Cross-correlation analysis using Pearson’s coefficient displayed as a heat map. (H,I) Zoom-in regions from panel (F) depicted in white rectangles, highlighting nanoscale structural detail and colocalization. Scale bar 2 µm.

Fiducial markers (gold nanoparticles) were incorporated throughout the experiment to enable accurate post-processing drift correction and precise overlay of sequentially imaged targets. System stability was quantified across multiple imaging and washing cycles: drift during 25-min imaging rounds was typically 100-300 nm, and washing steps introduced an additional drift of d200-500 nm. In contrast, manual buffer exchange often resulted in micrometer-scale displacements. Fiducial-based drift correction efficiently compensated for the observed drift, allowing for the accurate registration of all five imaging rounds without detectable misalignment. These results highlight the improved stability of automated fluidics compared to manual workflows.

Together, these results demonstrate that our microfluidics system enables robust multiplexed Exchange-PAINT imaging with high localization precision. The combination of efficient premixing labeling strategies, reproducible buffer handling, and fiducial-assisted drift correction establishes a workflow that is both practical and scalable for high-throughput SMLM experiments.

### Five-target multiplexed imaging of ventricular cardiomyocytes

To explore the applicability of our fluidics platform to more demanding biological systems, we performed Exchange-PAINT imaging of isolated mouse ventricular cardiomyocytes (CMs). To our knowledge, this represents the first demonstration of five-target super-resolution imaging in primary cardiomyocytes using DNA-PAINT. Multiplexed imaging of CMs poses unique technical challenges: these cells are large (∼100×30 µm) and fragile, requiring extremely gentle washing between imaging rounds to avoid detachment from the coverslip. Our microfluidics system proved particularly effective in this regard, performing thorough yet gentle washes without perturbing cell adhesion. This capability enabled the sequential imaging of multiple targets within the same cardiomyocyte. Imaging of multiple CMs confirmed the reproducibility of this workflow (Supplementary Figure S2).

The imaging plane was typically located several microns deep into the cell interior, resulting in reduced signal-to-background ratio compared with TIRF/HILO illumination in U2OS cells. Consequently, the average localization precision was approximately two-fold lower (23.9 nm, Table S1 in the SI). Drift correction was based on cross-correlation analysis, while a custom Python-based algorithm enabled accurate overlay of the sequentially acquired images.

Despite these challenges, microfluidics-enhanced Exchange-PAINT enabled new insights into the spatial organization and interplay of proteins expressed in specialized CM nanodomains, such as Caveolin-3 (CAV3), Ryanodine receptor type 2 (RyR2), Junctophilin-2 (JP2), Connexin-43 (CX43), and Dysferlin (DYSF) (see Figure 3 and CM schematic at the Supplementary Figure S2). By leveraging Exchange-PAINT on isolated ventricular myocytes, it becomes possible to simultaneously i) localize Caveolin-3 positive membrane domains called caveolae on the lateral surface sarcolemma and its regular membrane invaginations forming the transverse-axial tubule system^24^; ii) study the organization of sarcoplasmic reticulum RyR2 Ca^2+^ release channels clusters, iii) explore the local expression of the RyR2 binding and membrane-tethering protein Junctophilin-2 within the dyadic cleft and in close proximity to caveolae^25^; and iv) analyze the localization of Connexin-43 gap-junctional hemichannel plaques mediating the electrical coupling between CMs at the intercalated disc^26^. Interestingly, the subcellular expression of the Ca^2+^-sensitive membrane fusion and repair protein Dysferlin can be precisely mapped to all of these CM membrane nanodomains^27^ (Figure 3E), potentially stabilizing local membrane integrity and providing a basis for CM membrane plasticity. To validate the specificity of the Dysferlin signal, we performed control experiments in Dysferlin-knockout cardiomyocytes subjected to identical labeling and imaging workflows, which showed no Dysferlin signal while other protein targets were unaffected (Supplementary Figure S3). Cross-correlation analysis (Pearson’s coefficient) confirmed these structural relationships. CAV3, DYSF, and JPH2 displayed high colocalization, consistent with their cooperative function at membrane nanodomains^27^, while CX43 and RyR2 showed less overlap with these structures. The ability to resolve these membrane proteins within the same cell provides unprecedented detail regarding the nanodomain organization and function of CMs, shedding new light on key mechanisms in cardiac physiology and disease.

**Figure 3.**
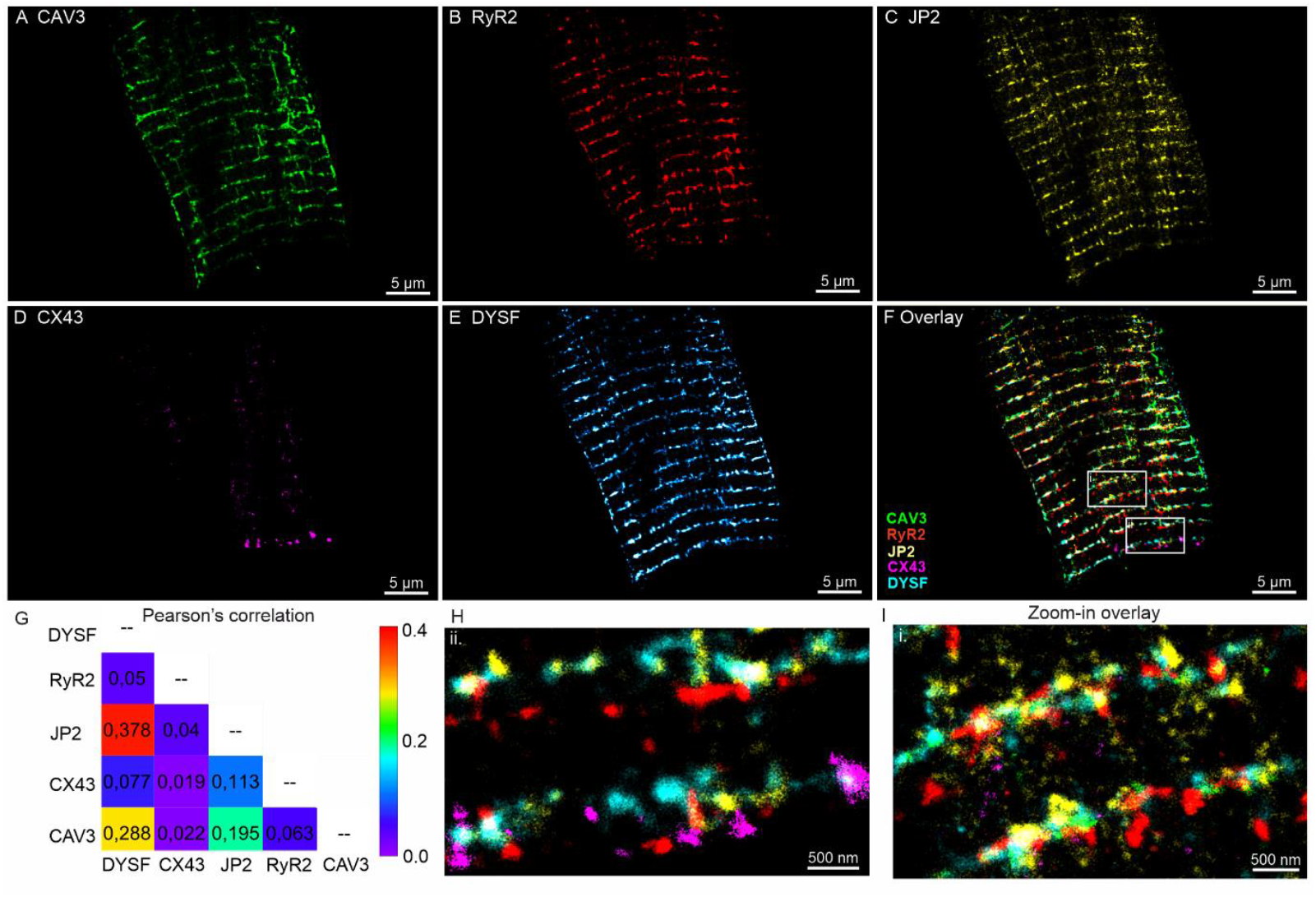
Microfluidics-enhanced Exchange-PAINT 5-target imaging of isolated ventricular cardiomyocytes. The following protein targets were imaged: (A) Caveolin-3 (CAV3); (B) Ryanodine receptor type 2 (RyR2); (C) Junctophilin-2 (JP2); (D) Connexin-43 (CX43); (E) Dysferlin (DYSF). (F) Overlayed image of all protein targets. (G) Cross-correlation analysis using Pearson’s coefficient displayed as a heat map. (H,I) Zoom-in regions of the overlay in panel (F), as indicated by white rectangles.

Together, these results establish fluidics-assisted multiplexed DNA-PAINT as a powerful approach for nanoscale imaging of primary cardiomyocytes. By overcoming the challenges of thorough washing, sample fragility, and deep imaging, our microfluidics system enabled the first 5-plex super-resolution imaging of isolated cardiomyocytes, providing detailed insight into the molecular architecture of cardiac nanodomains and their interplay in excitation-contraction coupling and membrane repair.

## Discussion

Multiplexed super-resolution microscopy is increasingly recognized as a standard tool in structural cell biology, yet manual Exchange-PAINT workflows remain labor-intensive, susceptible to mechanical sample drift, and difficult to standardize across different users. By integrating microfluidics, we provide precise and reproducible solution handling, fast exchange with minimal dead volume, and full automation of complex workflows. These capabilities are particularly important for fragile samples such as cardiomyocytes, where gentle solutions exchange prevents cells detachment from surface while thorough washing avoids cross-talk between cycles. This combination of factors enabled, for the first time, five-target Exchange-PAINT imaging of primary ventricular cardiomyocytes.

Beyond this application, the system’s adaptability extends its scope across microscopy platforms. Compatibility with confocal microscopes provides access to fluorescence lifetime– based multiplexing, while integration with spectral separation or advanced DNA-PAINT modalities such as RESI opens the way to hybrid strategies for robust, high-throughput nanoscale mapping. Coupling system automation with deep learning-based acquisition will enable adaptive workflows and reduce user-dependent variability.

Compared with commercial alternatives, our platform is cost-efficient, open-source, and straightforward to adapt to diverse configurations, including wide-field, confocal, STED, and MINFLUX. Its modular, stackable design ensures scalability without theoretical channel limits. Taken together, these features position microfluidics as a transformative technology of choice for multiplexed SRM, establishing a path toward standardized, high-throughput imaging for both fundamental research and translational diagnostics.

## Methods

### U2OS cell culture

We used genome-edited U2OS cell line Zyxin-rsEGFP2 (homozygous), introduced previously in Ref^28^ and U2OS-CRISPR-NUP96-mEGFP (Cytion #300174)^29^. The cells were cultivated in McCoys 5a medium (16600082, Thermo Fisher Scientific), supplemented with 100 U ml^−1^ penicillin, 100 ug ml^−1^ streptomycin (P0781, Sigma-Aldrich), 1 mM Na-pyruvate (S8636, Sigma-Aldrich) and 10% (v/v) FBS (FBS.S 0615, Bio and SELL) at 37^0^C, 5% CO_2_. The cells were cultured for 1 day on 35 mm glass bottom (No. 1.5) dishes (81158, Ibidi) and then fixed in pre-warmed 8% formaldehyde in PBS for 10 min. After washing with PBS pH 7.4 the dish was stored at 4^0^C before imaging.

### Sample preparation and labeling of U2OS cells

Unconjugated single-domain antibodies (sdAb, NanoTag Biotechnologies GmbH, Germany) carried a single ectopic cysteine at the C-terminus, allowing chemical coupling via thiol-reactive chemistry. DNA docking strands (Biomers GmbH, Germany) were functionalized with a 5′-azide group and conjugated to sdAb using a dibenzocyclooctyne (DBCO) crosslinker^30^. Immunolabelling was performed using a staining solution comprised of permeabilization buffer (PB), primary single-domain antibodies (1.sdAb) conjugated to DNA docking strands and premixtures of primary antibodies (1.Ab) and secondary single-domain antibodies (2.sdAb) conjugated to DNA docking strands^22^. For immunolabelling cells were incubated for 2h at RT under gentle shaking.

GFP-tagged zyxin was revealed using 25 nM of anti-GFP 1.sdAb (NanoTag Biotechnologies GmbH, Germany; Cat. No. N0305) coupled with the R4* concatenated DNA docking sequence (5′-ACACACACACACACACACA-3′). To reveal paxillin, we employed a premixing protocol^22^ where, recombinant anti-paxillin (pPax-Y118) 1.Ab (Abcam, ab32084, 15 nM) was incubated for 30 min with 2.sdAb anti-Rabbit IgG (NanoTag Biotechnologies GmbH, N2405; 45 nM) conjugated to the R1* docking DNA strand (5′-TCCTCCTCCTCCTCCTCCT-3′)^31^. Microtubules were labeled using an anti-α-Tubulin mouse IgG1 1.Ab (Synaptic Systems, Cat. No. 302211, Germany), premixed with 2.sdAb anti-Mouse IgG1 (NanoTag Biotechnologies GmbH, N2005) conjugated to the R3* docking DNA strand (5′-CTCTCTCTCTCTCTCTCTC-3′)^31^. Vimentin was labeled using an anti-Vimentin rabbit 1.Ab (Abcam, ab92547, Germany), premixed with 2.sdAb anti-Rabbit IgG (NanoTag Biotechnologies GmbH, N2405) coupled to the R6* docking DNA strand (5′-AACAACAACAACAACAACAA-3′)^31^, see detailed combinations in Table S3 in the SI.

After staining, cells were washed three times over 15 min, rinsed with PBS, and post-fixed with 4% paraformaldehyde (PFA) for 15 min at RT. Residual aldehydes were quenched with 0.1 M glycine in PBS, and samples were stored in PBS at 4 °C until imaging. To reveal actin, we used LifeAct conjugated to Cy3B (Cambridge Research Biochemicals, crb1130929h, UK). The stock solution was diluted to 1 µM, aliquoted, and stored at –20 °C. For imaging, LifeAct-Cy3B was used at ∼0.3 nM in PBS buffer (pH 7.4) supplemented with 500 mM NaCl.

### Imager strands

Fluorophore-labeled imager strands were purchased from Eurofins Genomics (Germany). Atto550-labeled imagers (full list in Supplementary Table S2) were dissolved in TE buffer (10 mM Tris, 1 mM EDTA, pH 8.0) at 1 µM, aliquoted, and stored at –20 °C. Prior to imaging, imagers were thawed and diluted to a final concentration of 0.2 nM in PBS buffer (pH 7.4) with 500 mM NaCl. Approximately 1 mL of imager solution was applied to the sample chamber for DNA-PAINT imaging.

### U2OS cells imaging and data analysis

Exchange-PAINT measurements were performed on a custom-built setup as described previously^32^ and in the Supplementary Information. Regions of interest (ROIs) of up to 50 × 50 µm^2^ were recorded with a pixel size in the object plane of 103.5 nm, an exposure time of 30 ms, and an emCCD gain of 500. Each dataset consisted of 50,000 frames. The room temperature was maintained at 23.0°C to ensure thermal stability of the setup. Linear sample drifts of typically 1-2 pixels were observed during acquisition and were corrected using fiducial markers. Excitation was performed in a highly inclined and laminated optical sheet (HILO) configuration with a laser power of ∼25 mW, providing optimal signal-to-noise ratios. Data analysis has been performed using free software FiJi^33^, plugin ThunderSTORM^34^. Identical data analysis workflow and parameters were applied as in previous work^32^.

### Cardiomyocytes

#### Mouse ventricular myocyte isolation

Animal procedures were conducted in accordance with the guidelines from Directive 2010/63/EU of the European Parliament on the protection of animals used for scientific purposes, and approved by the local veterinarian state authority (LAVES, Oldenburg, Germany; animal protocol no. T22.19). Adult sex-mixed mice in the C57BL/6J background were used for experiments. Mice were anesthetized with 2% isoflurane prior to cervical dislocation. Mouse hearts were rapidly excised and retrogradely perfused under a modified Langendorff-perfusion system for ventricular myocyte isolation using collagenase type II (Worthington)^27,35^.

#### Sample preparation and labeling of Cardiomyocytes

For multiplexed Exchange-PAINT imaging, isolated ventricular myocytes were fixed with 4% PFA in PBS on laminin-coated glass imaging dishes (Ibidi) for 10 minutes. Residual aldehydes were quenched with 0.1 M glycine in PBS for 15 min. Subsequently, cardiomyocytes were washed with PBS, permeabilized and blocked in PBS buffer containing 0.2% Triton X-100 and 10% bovine calf serum for 30 min at room temperature. The blocking buffer was supplemented with 0.1 mg/mL sheared salmon sperm DNA and 0.05% dextran sulfate^36,37^. Following blocking, Image-iT™ FX Signal Enhancer (Thermo Fisher Scientific) was applied for 10 min to reduce non-specific binding of imager strands, as described previously^36,37^. Nanobodies conjugated to DNA docking strands were diluted in staining buffer consisting of 3% bovine calf serum, 0.2% Triton X-100, and 0.05 mg/mL sheared salmon sperm DNA in PBS. This staining buffer enabled single-step labeling of cardiomyocytes. For each one-step immunofluorescence labeling, primary antibodies (1.Ab) were pre-mixed with secondary nanobodies conjugated to DNA docking strands at a molar ratio of 1:3^22^. The premixed antibody complexes were incubated for 30 min at room temperature with moderate orbital shaking, protected from light. To prevent crosstalk between different labeling complexes, a multiplexing blocker was added for 5 min. The Multiplexing Blocker Mouse (NanoTag, Cat No: K0102-50) was employed to saturate free binding sites on secondary nanobodies, thereby preventing unintentional cross-reactivity and ensuring specificity during multiplexed labeling^11^, full reagents list is provided in the SI, Table S4. After incubation with labeling complexes, cardiomyocytes were washed twice with staining buffer and subsequently incubated with Image-iT™ FX Signal Enhancer for 10 min at room temperature. Each primary Ab-Nb complex was initially prepared in a 20 µL PBS and incubated for 30 min at room temperature to ensure complex formation, see detailed combinations in Table S5 in the SI. Then, labeling complexes were diluted to the desired final concentration of a primary Ab in staining buffer and applied to the samples for overnight incubation at 4°C. After staining, cardiomyocytes were rinsed sequentially with staining buffer and PBS, and post-fixed with 4% PFA for 10 min at room temperature. Residual aldehydes were again quenched with 0.1 M glycine in PBS. Finally, cardiomyocytes dish was filled with PBS and stored at 4°C until further use. While handling cardiomyocytes samples, the number of washing steps in preparatory/labeling steps had to be minimized to avoid cells detachment from surface and keep the cardiomyocytes density high.

#### Cardiomyocyte imaging and analysis

First, the candidate ventricular cardiomyocyte has been selected. For this purpose, an increased concentration (∼2 nM) of imager R6 was introduced into the chamber. Once the suitable cell has been detected, the imaging plane was carefully selected. Then, the imager concentration was diluted by injection of the imaging buffer into the chamber, dropping the imager concentration to 0.05-0.1 nM. At this concentration, single molecule blinking events became visible. Then, the sample was gently flushed with the PBS buffer, until no signal has been visible. Subsequently, the next imager was injected and its concentration was adjusted to optimal conditions, where single blinking events could be identified. The data was recorded and the imaging cycle continued.

The following imaging parameters were used: white light laser was employed with excitation window 510-556 nm and power of ∼25 mW at the back of the objective. The emission filter 609/62 nm has been used to reject the excitation light. The following acquisition settings were used: camera gain 500, exposure time of 30 ms, total 50k frames were acquired. Identical data analysis has been used as for the U2OS cells and the previous work^32^.

## Data Availability

All data supporting the conclusions of this study are available from the corresponding authors upon reasonable request.

## Code availability

FiJi: https://fiji.sc/

Fiji ThunderSTORM plugin: https://github.com/zitmen/thunderstorm

## Notes

FO is a shareholder of NanoTag Biotechnologies GmbH. All other authors declare no competing interests.

## Supplementary information

Available. The following sections are included in the SI: Microfluidics system description; Manual and automated control of microfluidics system; Wide-field single molecule localization microscopy; Average localization precision for 5-plex SMLM imaging of U2OS and Cardiomyocytes; DNA-PAINT docking and imager sequences and modification details; U2OS cells labeling details; U2OS cell: schematics of imaged targets and imaging workflow; Ventricular Cardiomyocyte: schematics of imaged targets and imaging workflow; Cardiomyocyte handling reagents; Cardiomyocyte cells labeling; Dysferlin-knockout Cardiomyocyte imaging; Additional images of U2OS cells.

## Acknowledgment

The authors are grateful to Simon Bahl, Kevin Adner and Markus Schönekeß (DPI Electronics workshop) for the development of electronics for microfluidics operation. The authors thank Prof. Dr. Sven Thoms for supporting the project in the initial stage. The authors are grateful to Dr. Daniel Jans, Nicole Molitor and Prof. Dr. Stefan Jakobs for providing GFP-modified cell lines for Zyxin and NUP96, help with the sample preparation and supporting the project.

