## Supplementary Information for "A versatile microfluidics platform for enhanced multi-target super-resolution microscopy"

### Microfluidics system description

The platform operates using compressed air to pressurize reagent tubes, from which solutions are delivered to the experimental chamber. Pressures below 1 bar are sufficient for system operation. Compressed air can be supplied from laboratory infrastructure or a portable compressor. In our experiments, building-supplied air was split via T-connectors (QST-6, Festo) and 6 mm tubing (PUN-6x1-GE, Festo) and directed through a precision regulator (MS6, Festo), reducing pressure to 0–0.3 bar (0–4.4 psi) suitable for Exchange-PAINT workflows. When unfiltered air is used, an inline filter (PTA013, Thorlabs) is recommended.

The regulated air was routed to solenoid valves (MH1, Festo), which distributed pressure to selected channels in response to 24 V external control signals. Each solenoid output was connected to pressure tube holders (Fluiwell-4C for 2 mL tubes; Fluiwell-1C for 15 mL tubes, Fluigent) via 4 mm tubing (PUN-H-4X0.75, Festo). Holders were cleaned after each experiment, while new reagent tubes were used each time (2 mL: Micrewtube T341-6T, Simport; 15 mL: Greiner Bio-One™, Fisher Scientific). Imager solutions (0.2–0.5 nM in PBS with 500 mM NaCl) were supplied from 2 mL tubes, while larger 15 mL tubes were used for wash buffer (PBS).

To minimize dead volume, tube holders were mounted on a 25 mm construction rail (XE25L500/M, Thorlabs) positioned close to the sample chamber. They were aligned at chamber height using a post (P350/M, Thorlabs) and clamp (C1511/M, Thorlabs) to prevent gravity-driven flow. Additional components were mounted on an adapter plate (MB1111A/M, Thorlabs). These refinements improved upon our earlier prototype design[1].

Solutions were delivered to the chamber through biocompatible Tygon tubing (VERNAAD04103, VWR) connected to a transparent custom cover adapted to fit most standard open chambers (e.g., Ibidi Petri dishes or 8-well plates). The open format facilitated direct visual inspection of solution exchange, which was essential during development. To avoid undesired forward flow and backflow during fluids delivery, tubing should be filled completely and positioned at the same height as the sample chamber. Tygon tubing for solutions inlet/outlet shall be changed after each experiment to prevent salt accumulation, clogging and ensure smooth fluids delivery and removal. A simple inline air filter is recommended to prevent particulate contamination of the valves and to ensure smooth operation over time.

### Manual and automated control of microfluidics system

Control signals (24 V) for the solenoid valves were supplied either by a custom-built manual controller or by an automated system capable of addressing up to 48 channels and operated via LabVIEW software. In the manual configuration, an 8-channel switchboard directly triggered the valves for simple fluid channel selection. For automated operation, the controller consisted of a power supply, an Arduino-based microcontroller, and a signal-amplification stage that converted the Arduino’s 5 V logic output to the 24 V required by the solenoid valves. Each valve manifold contained eight valves; up to six manifolds (48 channels in total) could be connected to the controller through standard 15-pin cables. The LabVIEW graphical user interface (GUI) allowed users to define complex fluid-handling sequences-including channel order, timing, and duration-without additional coding, enabling intuitive programming of automated workflows.

Removal of solutions from the imaging chamber was performed either manually using syringes (5–20 mL) into 50 mL waste containers (CORN11706, VWR) or automatically using a peristaltic pump (MINIPULS 3, Gilson) triggered by TTL signals from the controller. The pump provided gentle and controlled fluid removal, which was especially advantageous for sensitive or extended washing steps.

**Wide-field single molecule localization microscopy**

*Optical setup description*

Wide-field measurements were performed using a custom-built optical setup, described elsewhere[1] and shown in Figure S1. A pulsed super-continuum white light laser (WL Laser) (SuperK Fianium, NKT Photonics) was used for sample excitation. A variable filter (VF) (SuperK Varia, NKT Photonics) connected to the laser output enabled flexible selection of the output wavelength (510-552 nm for efficient excitation of Atto 550 and Cy3B). A neutral density filter (NE10A-A, Thorlabs), in tandem with a variable neutral density filter (ND) (NDC-50C-4-A, Thorlabs), was used to adjust the laser excitation power.

The laser beam was coupled into a single-mode optical fiber (SMF) (P1-460B-FC-2, Thorlabs) with a typical coupling efficiency of 40%. After exiting the fiber, the collimated laser beam was expanded 3.6× using telescope lenses (TL1 and TL2). The typical excitation intensity at the sample was approximately 1.2 kW/cm² (24 mW at the entrance to microscope). The laser beam was focused onto the back focal plane of the TIRF objective (UAPON 100X oil, 1.49 NA, Olympus) using an achromatic lens (L1) (AC508-180-AB, Thorlabs). Beam displacement relative to the optical axis for switching between EPI, HILO, and TIRF illumination schemes was achieved using a translation stage (TS) (LNR25/M, Thorlabs). Smooth lateral positioning of the sample was enabled by a high-performance two-axis linear stage (M-406, Newport). In addition, an independent one-dimensional translation stage (LNR25/M, Thorlabs) combined with a differential micrometer screw (DRV3, Thorlabs) was used to move the objective along the optical axis for focusing.

Spectral separation of the collected fluorescence light from the excitation path was achieved using a multi-band dichroic mirror (DM) (Di03 R405/488/532/635, Semrock), which directed the fluorescence light toward the tube lens (L2) (AC254-200-A-ML, Thorlabs). The field of view was physically limited in the emission path by an adjustable slit aperture (SP60, OWIS) positioned in the image plane.

Lenses L3 (AC254-100-A, Thorlabs) and L4 (AC508-150-A-ML, Thorlabs) re-imaged the emitted fluorescence light from the slit onto an emCCD camera (iXon Ultra 897, Andor). Band-pass filters (BP) (BrightLine HC 692/40) were used to further block scattered excitation light. The total magnification of the optical system on the emCCD camera was 166.6×, resulting in an effective pixel size of 103.5 nm in the sample space. All experiments were conducted at 23 °C to ensure mechanical stability of the optical setup.

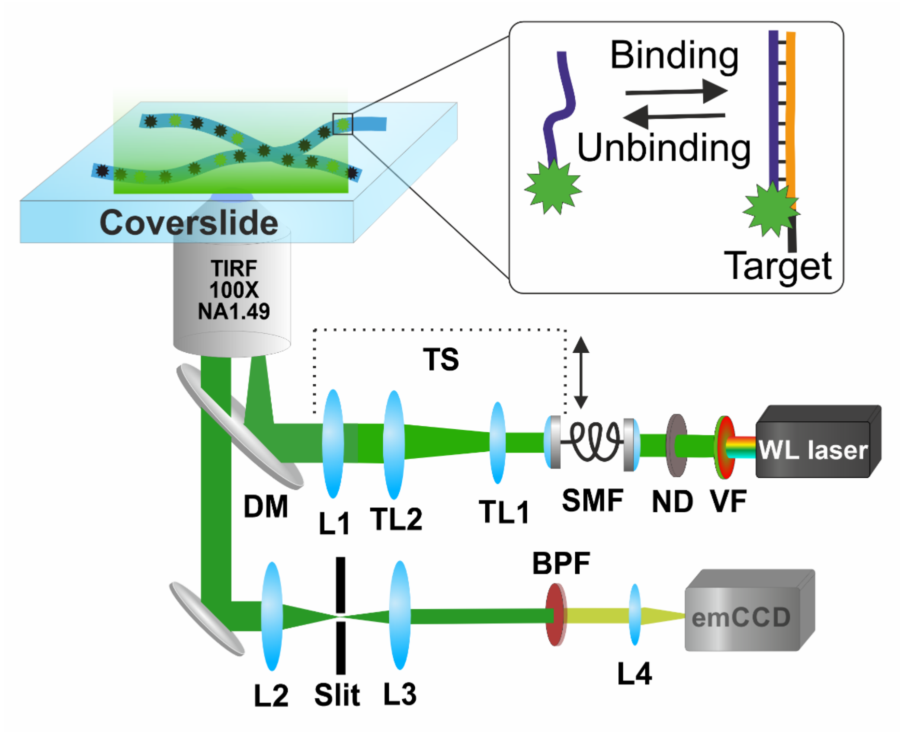


**Figure S1.** Schematics of home-built wide-field microscope used for multiplexed DNA-PAINT imaging. Available excitation schemes include EPI, HILO, and TIRF.

**Average localization precision for 5-plex SMLM imaging of U2OS and Cardiomyocytes**

| Target (U2OS) | Localization precision (nm) | Target (CM) | Localization precision (nm) |
|---|---|---|---|
| Microtubules | 10.3 | Caveolin-3 | 23.9 |
| Vimentin | 11.6 | Ryanodine receptor type 2 | 24.0 |
| Actin | 10.9 | Junctophilin-2 | 23.9 |
| Zyxin | 11.9 | Connexin-43 | 23.4 |
| Paxillin | 9.9 | Dysferlin | 24.1 |

**Table S1. Average localization precision for images in Figure 2 and 3 in the main text**

**DNA-PAINT docking and imager sequences and modification details.**

| DNA | Sequence (5’→3’) | Length (nt) | 5’ modification | 3’ modification |
|---|---|---|---|---|
| R1* | TCCTCCTCCTCCTCCTCCT | **19** | C3-Azide | --- |
| R2* | ACCACCACCACCACCACCA | **19** | C3-Azide | --- |
| R3* | CTCTCTCTCTCTCTCTCTC | **19** | C3-Azide | --- |
| R4* | ACACACACACACACACACA | **19** | C3-Azide | --- |
| R6* | AACAACAACAACAACAACAA | **21** | C3-Azide | --- |
| R6-Atto 488 | AACAACAACAACAACAACAA | **21** | C3-Azide | Atto 488 |
| R1 | AGGAGGATT | **9** | Atto 550 | --- |
| R2 | TGGTGGTTT | **9** | Atto 550 | --- |
| R3 | GAGAGAGAAA | **10** | Atto 550 | --- |
| R4 | TGTGTGTTT | **9** | Atto 550 | --- |
| R6 | TTGTTGTTT | **9** | Atto 550 | --- |

**Table S2. DNA PAINT docking and imager sequences and modifications.**

**U2OS cells labeling details**

| Target | Primary Ab | Secondary Nb | DNA Docking |
|---|---|---|---|
| Zyxin-GFP | ---- | sdAb anti-GFP, NanoTag, Cat. No: N0305 | R4* |
| Paxillin | Anti-Paxillin rabbit (pPax-Y118) antibody, Abcam, ab32084 | sdAb NanoTag N2405 (anti-rabbit) | R1* |
| Microtubules | anti-α-Tubulin mouse IgG1, Synaptic Systems, Cat. No: 302211 | sdAb NanoTag N2005 (anti-mouse) | R3* |
| Vimentin | anti-Vimentin rabbit antibody, Abcam, ab92547 | sdAb NanoTag N2405 (anti-rabbit) | R6* |
| PMP70 | anti-PMP70 rabbit IgG, Abcam, ab85550 | sdAb NanoTag N2405 (anti-rabbit) | R3* |
| NUP96-GFP | ---- | sdAb anti-GFP, NanoTag, Cat. No: N0305 | R4* |

**Table S3. U2OS cell targets labeling details.**

**U2OS cell: schematics of imaged targets and imaging workflow**

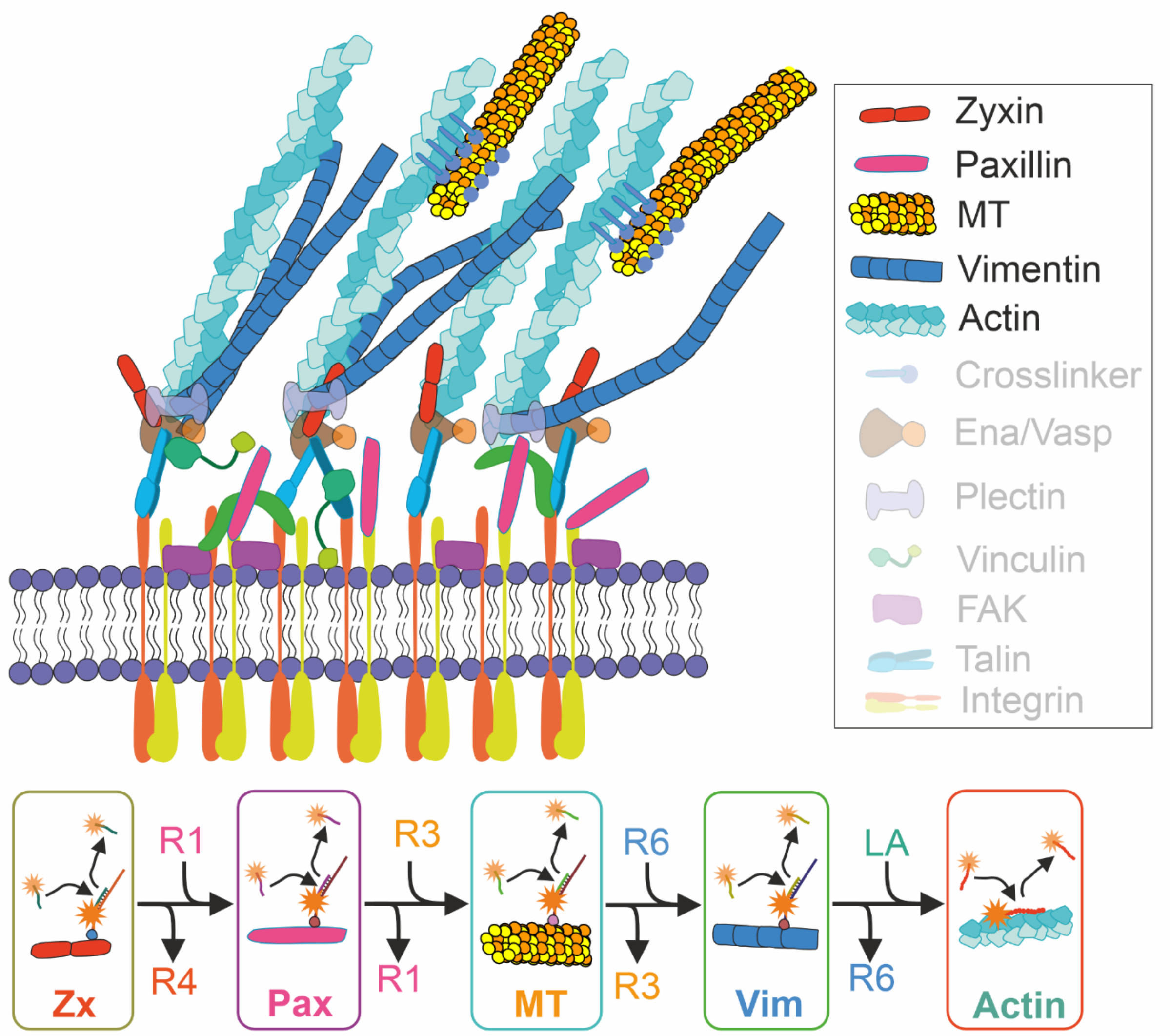


**Figure S1. Cellular targets for U2OS labeled with primary antibodies and functionalized secondary nanobodies.** Upper panel – cell bottom cut schematics with FA and cytoskeleton targets shown. Lower panel – Exchange-PAINT imaging workflow of U2OS cellular targets. R1-R4 and R6 imagers, as well as LifeAct were used to image the targets labeled with the complementary docking strands.

**Ventricular Cardiomyocyte: schematics of imaged targets and imaging workflow**

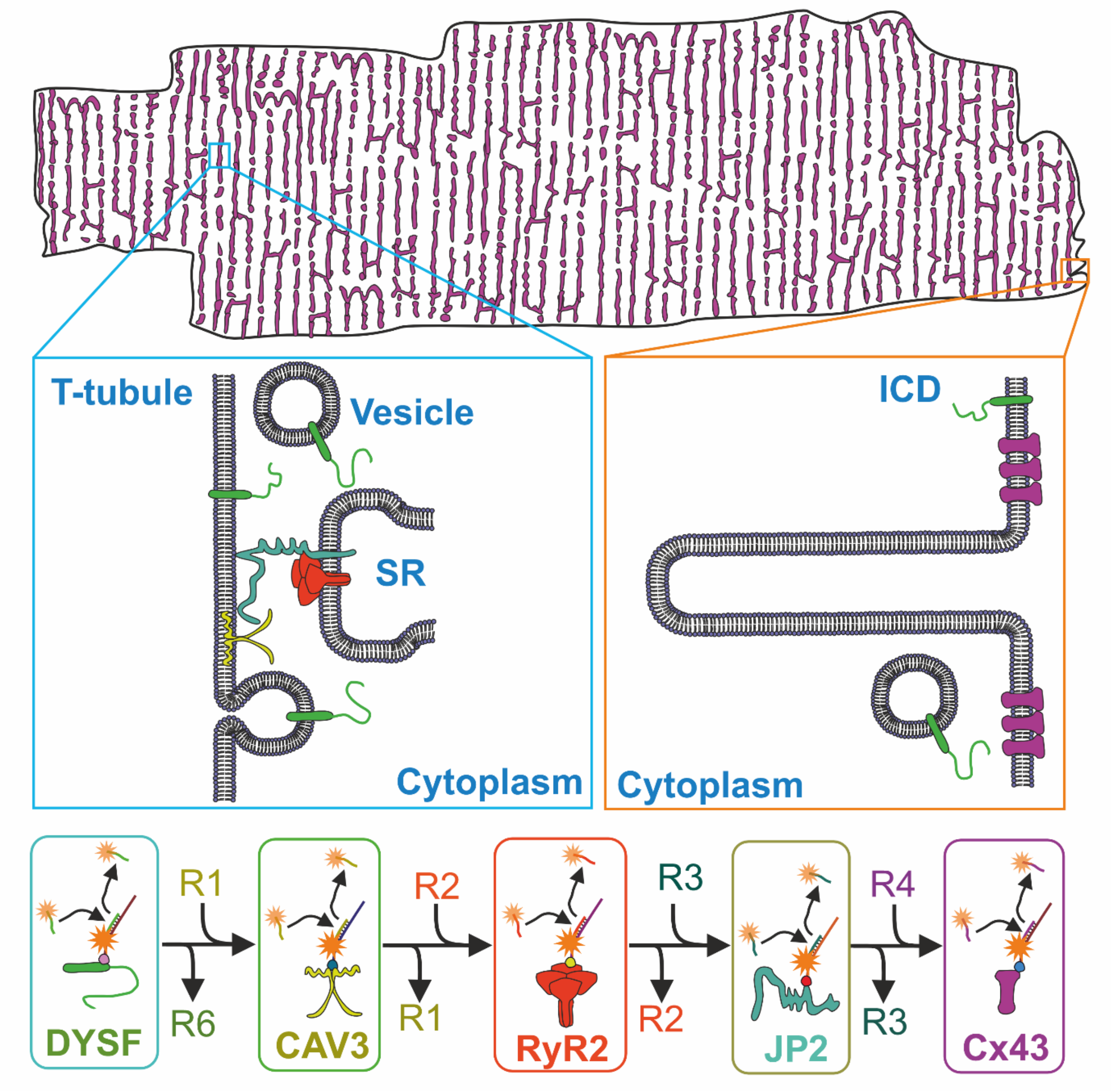


**Figure S2. Cardiomyocytes targets labeled with primary antibodies and secondary nanobodies.** Upper panel – whole ventricular cardiomyocyte cell. Zoom-ins show the arrangement of CM targets selected for imaging. Lower panel – Exchange-PAINT imaging workflow. R1-R4 and R6 are the imagers used to reveal the targets labeled with the complementary docking strands.

**Cardiomyocyte handling reagents**

| Reagent | Company | Cat. Number |
|---|---|---|
| sheared salmon sperm DNA | Thermo Fisher Scientific | 15632011 |
| dextran sulfate | Merck | D4911 |
| Image-iT™ FX Signal Enhancer | Thermo Fisher Scientific | I36933 |
| multiplexing blocker mouse | NanoTag | K0102-50 |

**Table S4. Cardiomyocytes handling reagents and materials.**

### Cardiomyocyte cells labeling

In the table below we provide the necessary details regarding the primary antibodies and secondary nanobodies used for labeling of five cardiomyocytes targets shown in Figure 3: Dysferlin, Ryanodine type 2 receptor, Caveolin-3, Junctophilin-2, Connexin-43

| Target | Primary Ab | Secondary Nb | DNA Docking |
|---|---|---|---|
| Dysferlin | Ab rabbit, IgG Cat. #ab124684, | sdAb NanoTag N2405 (anti-rabbit) | R6*- Atto 488 |
| Ryanodine type 2 receptor (RyR2) | Ab mouse IgG1, IgG1, Thermo Fisher Scientific, MA3-916, | sdAb NanoTag N2005 (anti-mouse) | R2* |
| Caveolin-3 | Ab mouse IgG1, BD Biosciences, Cat. # 610421 | sdAb NanoTag N2005 (anti-mouse) | R1* |
| Junctophilin-2 | Ab mouse IgG1, Santa-Cruz, Cat. # sc-398125 | sdAb NanoTag N2005 (anti-mouse) | R3* |
| Connexin-43 | Ab mouse IgG1, Cat. #610062 | sdAb NanoTag N2005 (anti-mouse) | R4* |

**Table S5. Cardiomyocytes targets labeling.**

### Dysferlin-knockout Cardiomyocyte imaging

Unless otherwise indicated, adult male and female C57BL/6J mice aged 8–16 weeks were used for all experiments. The dysferlin-knockout (KO) model was utilized as previously described.[2] Homozygous KO mice and wild-type (WT) littermate controls were generated by interbreeding heterozygous animals. Genomic DNA was isolated from ear punches, and genotyping was performed by polymerase chain reaction (PCR) using the following primer sets: for the WT allele, 5′-GCCAGACAAGCAAGGTTAGTGTGG-3′ and 5′-GCGGGCTCTCAGGCACAGTATCTGC-3′, yielding a 3400 bp product; for the KO allele, 5′-GCCAGACAAGCAAGGTTAGTGTGG-3′ and 5′-GCTGACTCTAGAGCTTGCGGAACC-3′, yielding a 3000 bp product.

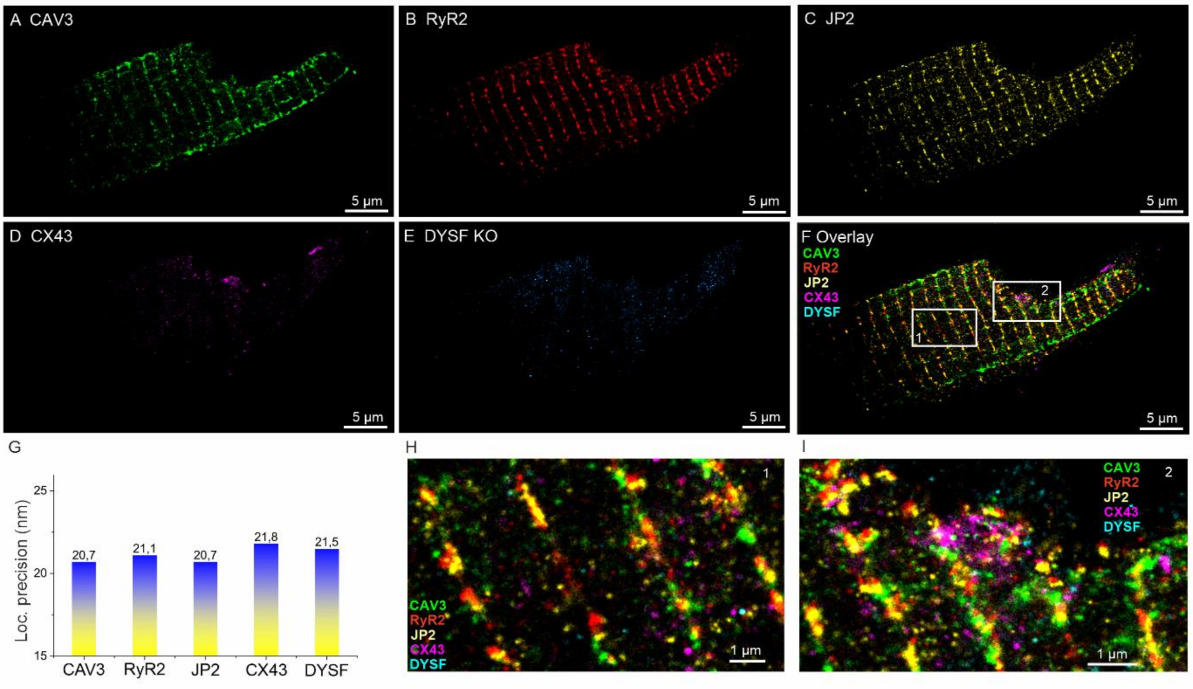


**Figure S3. 5-target microfluidics-enhanced Exchange-PAINT of Dysferlin knockout Cardiomyocyte.** Identical imaging protocol as for the CM cell in Figure 3 has been used. Targets: (A) Caveolin-3 (CAV3); (B) Ryanodine receptor type 2 (RyR2); (C) Junctophilin-2 (JP2); (D) Connexin-43 (CX43); (E) Knockout Dysferlin (DYSF). (F) Overlayed image of all protein targets. (G) Average localization precision of each imaged target. (H,I) Zoom-in views of the overlay in panel (F), as indicated by white rectangles.

### Additional images of U2OS cells

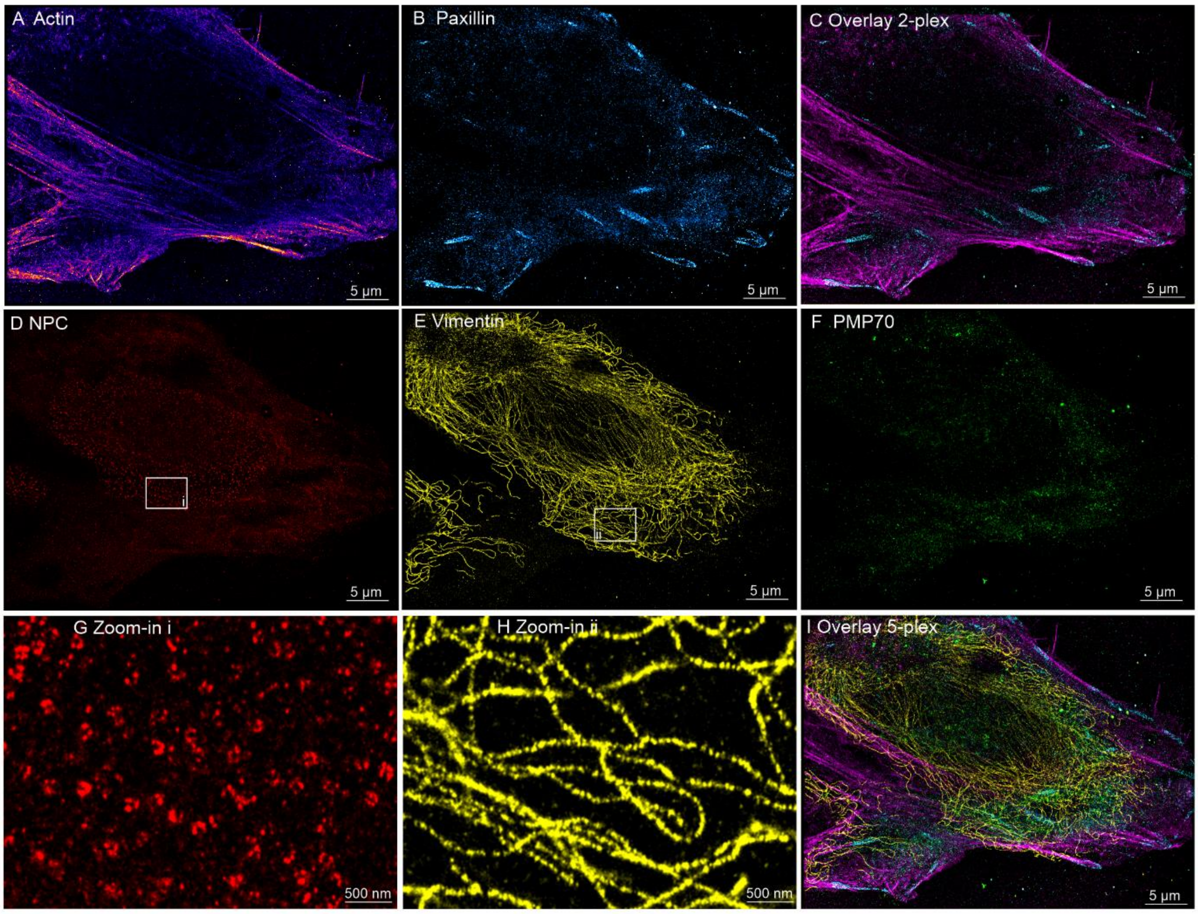


**Figure S4. Microfluidics-enhanced Exchange-PAINT 5-target imaging of U2OS cell #2.** Sequential imaging of U2OS cell targets: (A) Actin, (B) Paxillin, (C) Overlay of images in (A) and (B), (D) Nuclear pore complex (NPC) – NUP96, (E) Vimentin, (F) Peroxisomes (PMP70), (G,H) Zoom-in regions from panels (D,E) depicted in white rectangles, (I) Composite overlay of all channels – 5-plex image.

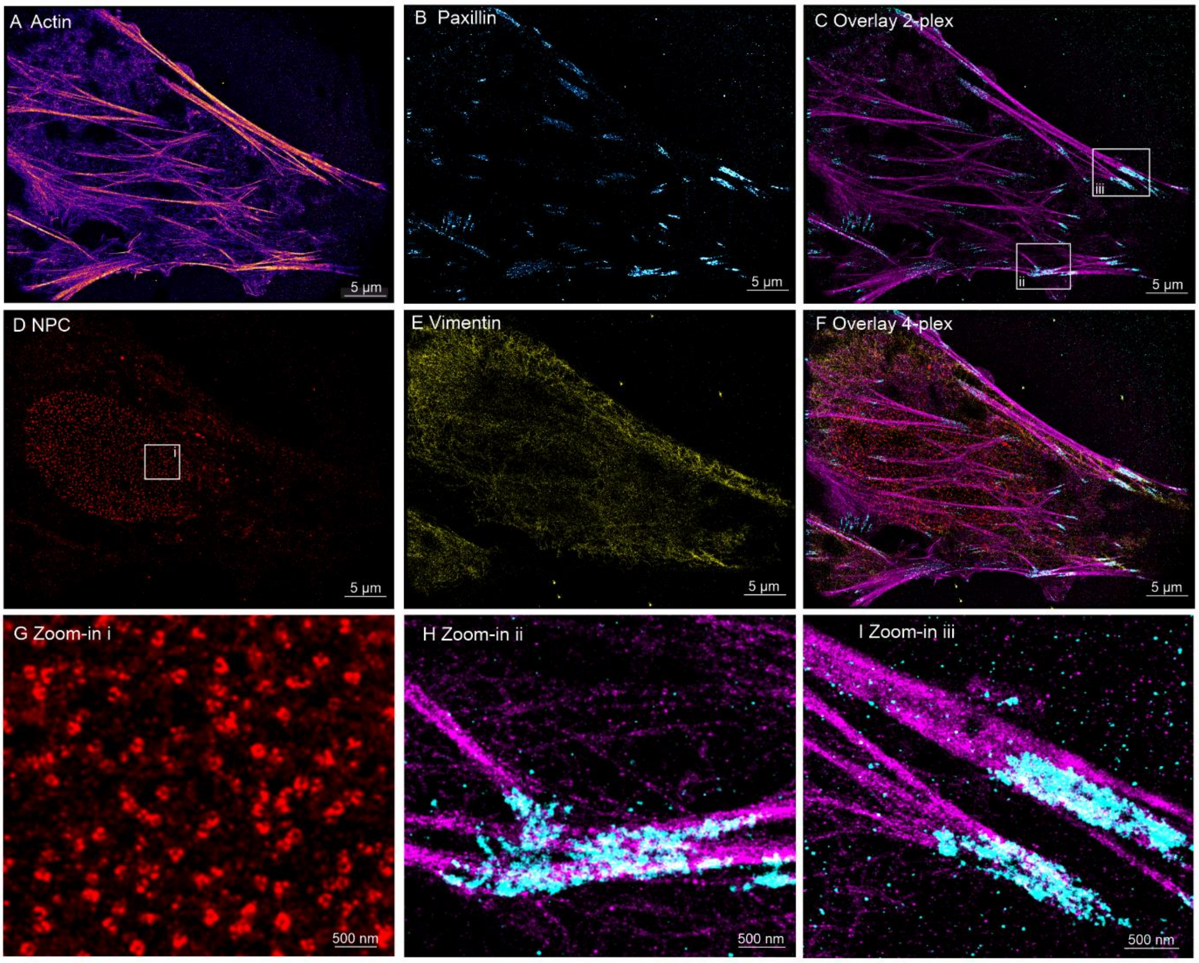


**Figure S5. Microfluidics-enhanced Exchange-PAINT 4-target imaging of U2OS cell #3.** Sequential imaging of U2OS cell targets: (A) Actin, (B) Paxillin, (C) Overlay of images in (A) and (B), (D) Nuclear pore complex (NPC) – NUP96, (E) Vimentin, (F) Composite overlay of all channels – 5-plex image. (G,H,I) Zoom-in regions from panels (C,D) depicted in white rectangles.
